# Illuminating the Potential: Light Harvesting and Semiconducting Properties of Bacterial Microcompartment Shell Proteins

**DOI:** 10.1101/2024.12.16.628711

**Authors:** Silky Bedi, S M Rose, Sharmistha Sinha

## Abstract

Biomaterials with self-assembly, genetic tunability, self-healing, and biocompatibility hold promise for next-generation electronics. This study explores disc-like shell proteins from bacterial microcompartments as photovoltaic materials. These self-assembling proteins exhibit semiconducting properties, including a low work function and non-linear I-V behavior. Notably, under UV light, they generate significant photocurrent without external voltage, demonstrating efficient electron transfer. Electron conduction occurs via tunneling, enabled by distinct electron-rich surface regions. High responsivity and quantum efficiency outperform prior protein-based systems, emphasizing their potential in photocurrent and light-harvesting applications. Genetic mutations revealed a proton-coupled electron transfer (PCET) mechanism, where tyrosine residues strategically positioned near proton abstraction sites enhance photocurrent generation. This study highlights the innovative capabilities of shell proteins for energy-efficient devices and demonstrates how genetic engineering can further optimize biomaterial performance, paving the way for sustainable electronic applications.

---

Modern-day technology is increasingly focused on developing devices that are autonomous or powered by renewable energy sources^1^. This shift is driving advancement in biomolecular electronics, with particular emphasis on the use of biomolecules for wearable technologies^2^, organ-on-a-chip models^3^, and medical diagnostics. In this evolving landscape, self-assembled peptides, proteins, and hybrid organic-inorganic protein conjugates are emerging as key players in bioelectronics ^4,5^

Their natural biocompatibility, self-assembly, self-healing capabilities, and ease of genetic tunability make them ideal candidates for next-generation technological solutions^6–8^. Proteins are particularly significant due to their role in biological energy conversion processes, such as respiration and photosynthesis, where they facilitate essential electron transport^9–11^. Certain amino acids within proteins act as electron reservoirs or transporters, and their proximity to electron acceptors enables the formation of efficient electron transport chains^12–15^. In some cases, the presence of aromatic amino acids like tyrosine and tryptophan, in specific environments and conformations, can absorb light to excite and mobilize electrons, imparting intrinsic conductivity to proteins^16–19^. This opens up exciting possibilities, especially with certain classes of proteins, for developing innovative bioelectronic devices^7^. The precise tiling of these proteins results in extended, planar mats **(Figure 1ai-iii)** that are remarkably stable and capable of spanning large areas, forming the protective exterior shell of the BMCs^20,21^. This organized arrangement not only reinforces the structural integrity of the BMCs but also plays a critical role in their function, particularly as selectively permeable barriers^22^.

**Figure 1:**
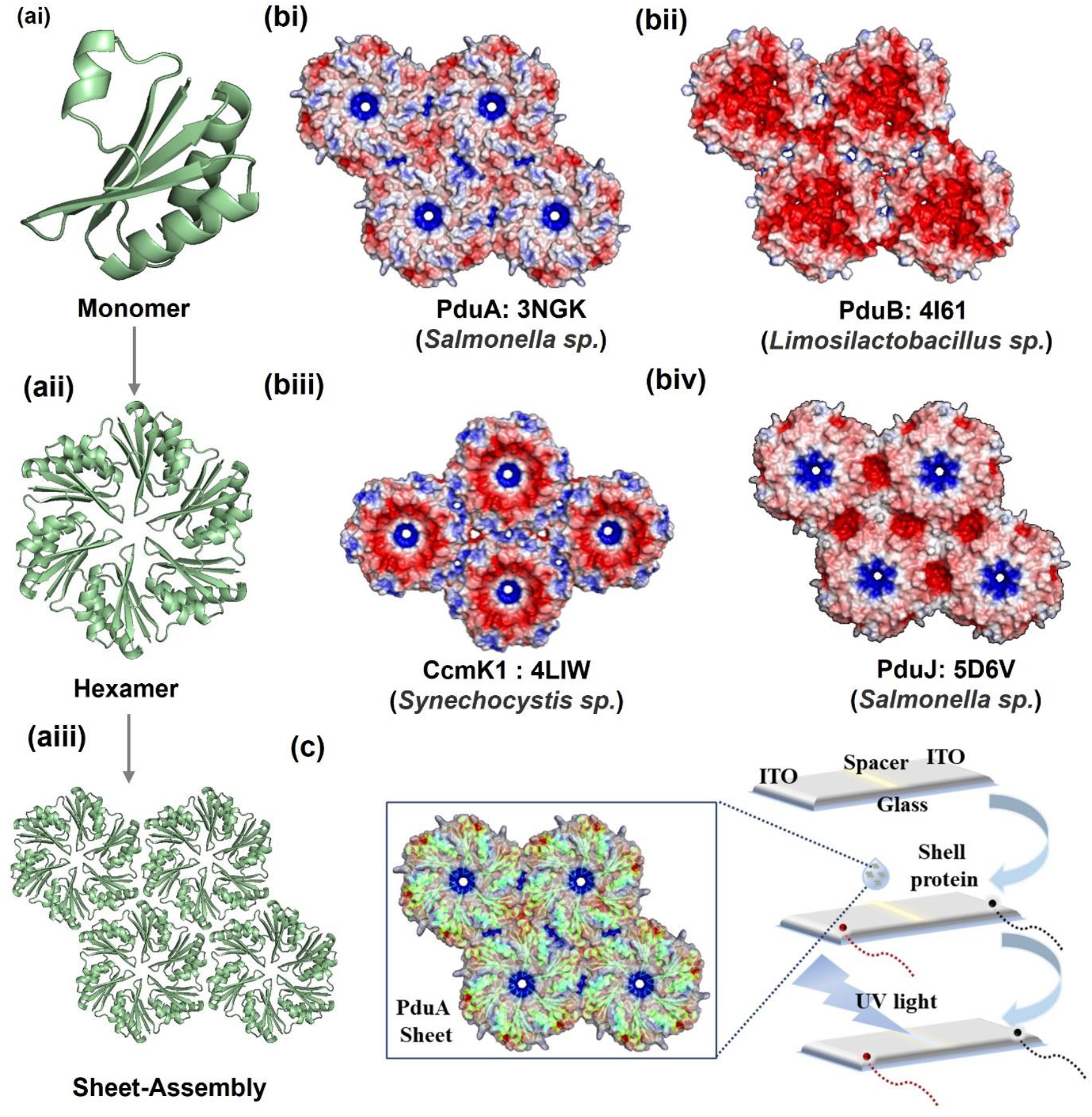
Asymmetric electron density on shell proteins facilitating electron conduction: (a-i) Monomeric unit of PduA (PDB ID: 3NGK) that assembles to form hexamers (aii). Hexamers undergo edge to edge association to form flat sheet assemblies (aii); (b-i) Electron density map of PduA (PDB ID: 3NGK); (b-ii) PduB (PDB ID: 4I61); (b-iii) β-carboxysome shell protein CcmK1, structurally analogous to PduA shell protein of PduBMC (PDB ID: 4LIW); (b-iv) PduJ shell protein from *Salmonella* species (PDB ID: 5D6V). These images illustrate distinct electron density patterns on the protein surface facilitating electron conduction over the protein surface; (c) Sheet assembly model of the PduA shell protein overlaid with its electron density pattern, highlighting potential electron conduction regions and its fabrication on the indium tin oxide (ITO) substrate for conductivity experiments.

The bacterial microcompartment shell proteins across several species exhibit discrete electron density patterns across the protein surface **(Figure 1 bi-iv)**. The electron density on both sides of the disc shows distinct patterns, and in some cases is reported to influence the selective permeability of the shell protein toward different substrates^23,24^. However, these electron density patterns result in localized electron-rich or electron-deficient regions on the shell protein surface, which may create potential differences conducive to the electron transfer across the shell protein surface.

In this work, we explored the asymmetry in electron density to study electron flow across these protein sheets. We focused on the major shell proteins PduA and PduBB’ from *Salmonella* PduBMCs, investigating their photo-assisted electron transfer capabilities using an in-house setup shown in **Figure 1c**. Given the abundance of tyrosine residues in these proteins **(Table S1)**, we hypothesized that their spatial arrangement within these shell proteins might facilitate electron transfer mobilization upon photoirradiation. Additionally, the self-assembly nature of these proteins enables the clustering of tyrosine residues, potentially amplifying electron flow and enhancing their utility in bioelectronics. By exploiting these precise tiling and electron density patterns, these protein mats could be used to create well-ordered networks for electron conduction, paving the way for innovative bioelectronics applications.

PduA forms flat molecular two-dimensional sheets upon overexpression in a bacterial host and further purification. Electron micrographs of PduA obtained using Transmission Electron Microscopy (TEM) and Field Emission Scanning Electron Microscopy (FESEM) show extended sheets **(Figure 2 ai-iv)**. We first examined the electron conduction properties of the PduA sheet by recording their I-V characteristics. The PduA protein is drop-casted onto a spacer created on an ITO substrate, as shown in **Figure 1c**. This setup bridges the connection between the ends of the substrate, which are then linked with copper wires to record current flow. The recorded I-V characteristics exhibit a non-linear response when subjected to a ±7 V bias voltage, **(Figure 2b)**. This non-linear behaviour provides strong evidence of the semiconducting nature of the PduA protein. To check the photoresponsivity of the shell protein, we first determined the work function value for PduA using Ultraviolet Photoelectron Spectroscopy (UPS). The work function indicates the minimum energy required to eject an electron from the surface of a material^25^. PduA protein sheets are drop casted on the surface of ITO. Complete coverage of the protein on ITO surface is confirmed by Coomassie Brilliant Blue G250 dye staining **(Figure S1a)**. The calculated work function for PduA is 2.96 eV, placing it within the typical range of semi-conducting materials^26^ **(Figure 2c)**. The control ITO shows a higher work function of 4.23 eV **(Figure S1b**) which is in agreement to the value reported in the literature^27^.The I-V curve for the bare ITO substrate shows a linear response, confirming its conductive behaviour, while that with empty channel casted shows no conductivity **(Figure S1c-d)**. The protein sample when drop casted in the channel bridges the gap, enabling electron flow across the ITO substrate, as demonstrated by the change in the I-V curve **(Figure S1e)**.

**Figure 2:**
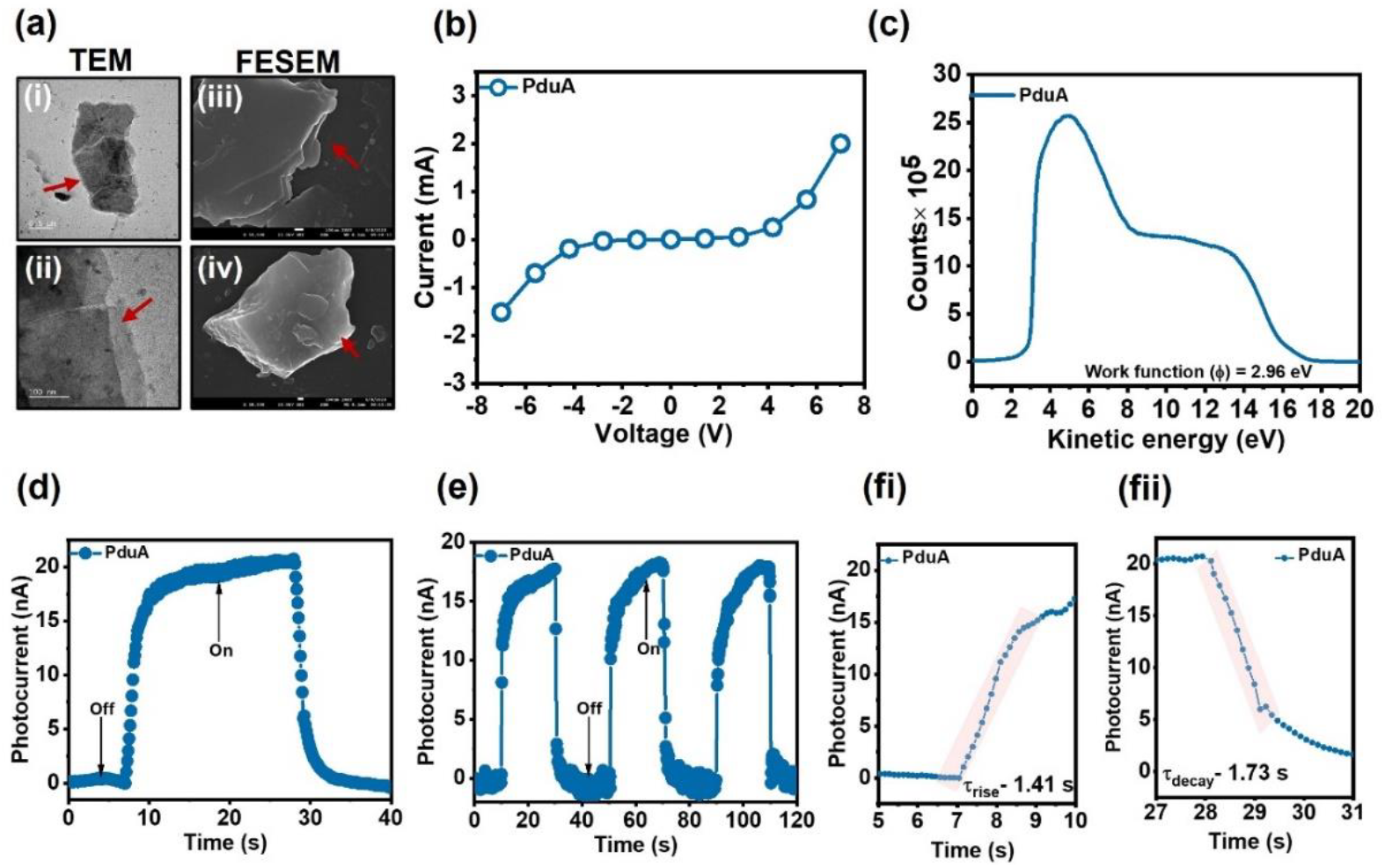
Electron conductivity and photocurrent generation in PduA shell protein: (a) TEM and FESEM imaging of PduA shell protein, illustrates sheet morphology of the protein (b) I-V characteristics of PduA, showing non-linear behavior, indicative of its semiconducting nature; (c) The measured work function of PduA, which falls within the typical range for semiconducting materials, indicating the ease of electron ejection from the surface of the PduA sheet; (d) Photocurrent generation in PduA-fabricated samples under UV light illumination (254 nm) without the application of external bias, demonstrating the intrinsic light-harvesting capability of the PduA shell protein; (e) On-off cycles of photocurrent response recorded during alternating dark and UV light exposure, showing consistent and reversible photocurrent generation; (f) Area used to calculate the response times for photocurrent rise τ_rise_ and τ_decay_, highlighting the fast and stable response of PduA to UV light, confirming its potential for photocurrent applications.

We then exposed the PduA sample to UV light (254 nm) and measured current generation without any externally applied voltage **(Figure 2d)**. The photo-response of PduA under dark and UV light illumination (at a power of ∼0.11mW) shows a current with amplitude of ∼20 nA. Repeated light on-and-off cycles for photocurrent generation for over 120 s demonstrated stable photocurrent generation consistent throughout the cycles **(Figure 2e)**.

We compared the performance of the shell proteins with globular protein, BSA. Under a ±7V bias, BSA produced negligible current (0.5 mA) **(Figure S2a)**. To investigate the photoresponsive properties further, we calculated the work function of BSA. The higher work function of BSA (3.2 eV) resulted in reduced photocurrent generation under UV light exposure, as shown in **(Figures S2b and S2c)**. This comparison suggests that the PduA sheet formed of laterally interacting hexagonal discs of PduA shell protein is more conducive to electron transport than globular proteins.

We also measured the rise and decay times (τ_rise_ and τ_decay_) to evaluate PduA’s response to light power. A shorter rise time is critical for applications requiring fast detection of light signals. PduA exhibited a τ_rise_ of 1.41 s and a τ_decay_ of 1.73 s, reflecting its quick photoresponsivity to UV illumination **(Figure 2fi-ii)**. These findings underscore PduA shell protein potential in photodetector applications. To gain a deeper understanding of the photoresponsivity of PduA, we next examined the mechanism behind the photocurrent generation in PduA.

To investigate the underlying mechanisms contributing to the current behavior of the shell proteins, we conducted I-V measurements of the PduA protein at two temperatures: 5°C and 25°C **(Figure 3a)**. We observed no significant change in current response across these temperatures, suggesting that conduction in shell proteins primarily involves electron tunneling^28^. To further, probe the mechanism behind photocurrent generation, we draw inspiration from natural light-harvesting systems in photosynthesis^11,29^. In photosystem II (PSII), chlorophyll (P680) absorbs light, resulting in a chlorophyll radical (P680^+^). The presence of aromatic amino acids in the photosynthetic system facilitates the Proton-Coupled Electron Transfer (PCET) reaction. In this process, both a proton and electron are transferred, either simultaneously or stepwise^30^. Within the photosynthetic reaction centre system, electrons generated from the water-splitting cluster (CaMn_4_) are transferred to the P680^+^ radical through the redox-active tyrosine residue, tyrosine (Yz)^31^. This electron transfer is coupled with transferring a phenolic proton from Yz to a nearby histidine residue (HisZ), which abstracts the proton by forming a hydrogen bond with Yz enabling the efficient electron and proton transfer.

**Figure 3:**
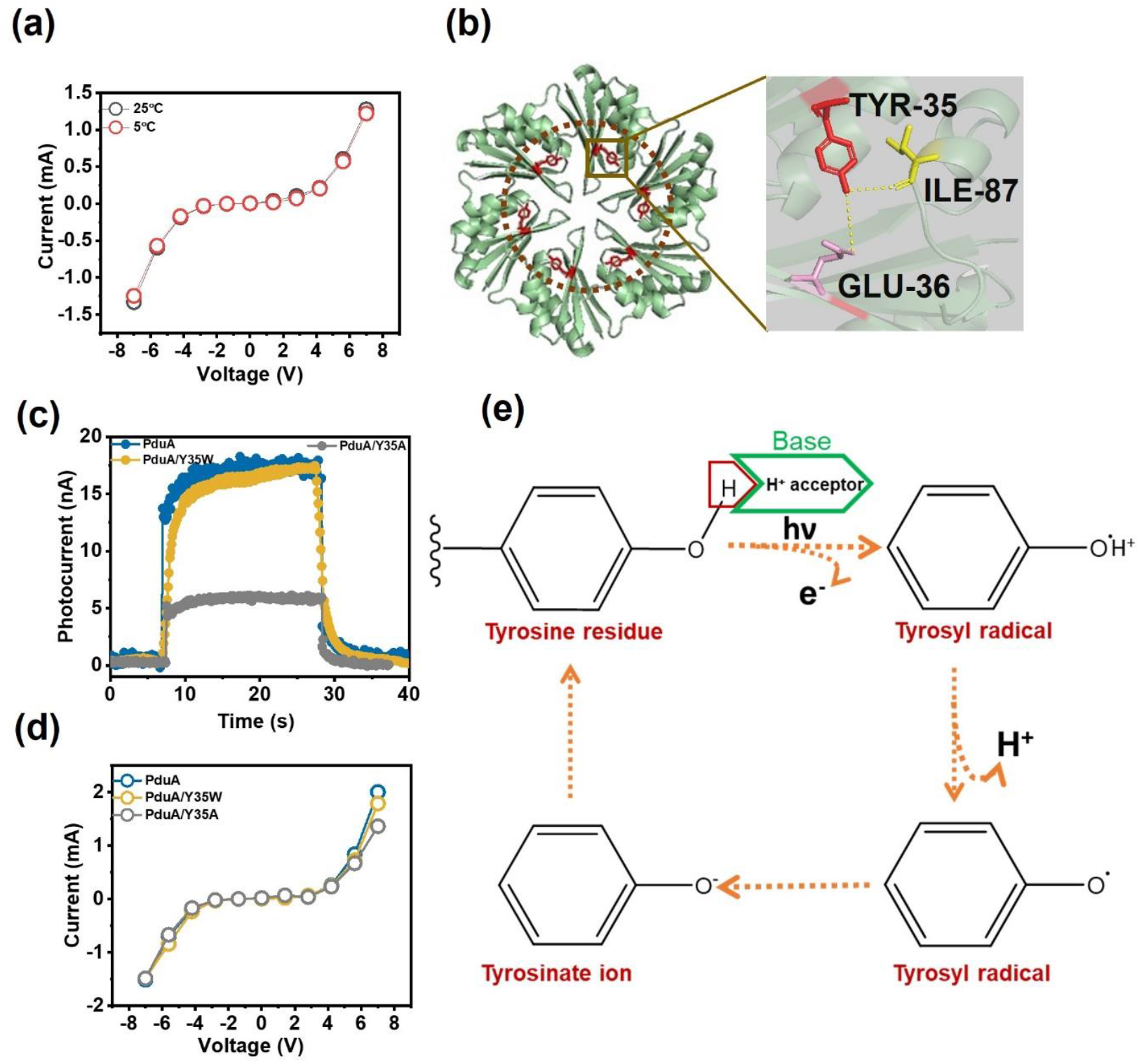
Mechanism of tyrosine-induced protein-coupled electron transfer (PCET) in PduA shell protein: (a) temperature-dependent photocurrent generation in PduA representing electron tunneling as conduction mechanism; (b) Spatial arrangement of tyrosine residues within the PduA hexamer, illustrating the proximity between these residues, which facilitates efficient electron transfer across the protein surface; (c) Photocurrent generation comparison, indicating a decrease in photocurrent when tyrosine is mutated to alanine (PduA Y35A), while the replacement of tyrosine with tryptophan (PduA Y35W) does not significantly impact photocurrent values, suggesting the importance of aromatic residue presence for maintaining photocurrent efficiency. e; (d) I-V curves for wild-type PduAWT and its mutants PduA Y35A (tyrosine to alanine) and PduA Y35W (tyrosine to tryptophan), showing no significant alteration in the semiconducting properties of the shell protein across mutations; (e) Schematic of PCET process in PduA shell protein, showing the formation of tyrosyl radical from tyrosine residue. The glutamate residues involved in hydrogen bonding with tyrosine participate in electron abstraction, initiating the PCET cycle. This mechanism highlights the role of tyrosine in enabling electron transfer within the PduA shell protein.

Inspired by this mechanism, we investigated tyrosine-mediated PCET in the self-assembled PduA wild-type sheet. PduA has a tyrosine residue at the 35^th^ position (Y35). The spatial arrangement of six Y35 residues within the PduA hexamer is shown in **(Figure 3b)**, where the hydroxyl group of the tyrosine residue form hydrogen bonds with adjacent glutamate (E36) and isoleucine (I87) residues. To assess the role of Y35 in photocurrent generation, we created a mutant, PduA[Y35A], replacing tyrosine at 35^th^ position with alanine **(Table S2)**. This mutation significantly reduced the photocurrent response compared to PduA WT **(Figure 3c)**, underscoring the importance of Y35. However, when Y35 is substituted with tryptophan PduA [Y35W] **(Table S3)**, there is no significant change in either the semiconductive properties **(Figure 3d)** or photocurrent response **(Figure 3c)**, suggesting that both tyrosine and tryptophan contribute to PCET in photocurrent generation **(Figure 3e)**^32,33^.

After confirming tyrosine’s involvement in the electron transfer, we examined potential hydrogen bond acceptors in the PduA hexamer. Using the Ligand-Protein Contacts & Contacts of Structural Units (LPCCSU) server^34^, we identified glutamate (E36) and isoleucine (I87) as possible acceptors of the phenolic hydrogen from tyrosine 35 in PduA. Glutamate is reported to participate in PCET in ribonucleotide reductase^35^. Glutamate-36 in PduA forms a hydrogen bond with Y35 through its side chain, while isoleucine makes the bond via its backbone carboxyl oxygen, as depicted in **Figure 3b**. However, proton acceptance by I87 could destabilize the protein’s structure. In conclusion, we propose that Y35 in PduA generates electrons under UV light (254 nm) through a simultaneous proton transfer to the adjacent E36 residue. Hence, tyrosine-mediated protein-coupled electron transfer appears to drive photocurrent generation in PduA. This led us to hypothesize that increasing the number of tyrosine residues within the shell protein sheet structure could further enhance photocurrent output. Building on this, we conducted experiments with a shell protein containing three tyrosine residues per monomer, resulting in nine tyrosine residues in the PduBB’ trimer ^36^.

PduBB’ self-assembles into flat sheet structures, as shown by TEM and FESEM images illustrating a sheet morphology similar to that of PduA **(Figure S4a)**. The electron density map of PduBB’ reveals high-density regions on its surface **(Figure 1b-ii)**, which likely contribute to enhanced electron conduction **(Figure S4b)**.

We assessed the electron conduction properties of PduBB’ by recording its I-V characteristics under a ±7 V external bias. PduBB’ exhibited significantly higher current flow than PduA shell protein **(Figure 2b, Figure S4b)**. This observation prompted an investigation into the effect of the increased number of tyrosine residues on the photoresponsivity of the shell protein. The work function of PduBB’ is determined to be 2.68 eV which is lower than that of PduA **(Figure S4c)**. This lower work function aligns with the observed photoresponse curve, indicating that PduBB’ generates three times more photocurrent than PduA **(Figures S4d and 2d)**. These findings support our hypothesis that a higher number of tyrosine residues enhances photocurrent generation, reinforcing the proposed mechanism for photocurrent generation in shell proteins. Moreover, PduBB’ demonstrated consistent, high photocurrent generation across multiple on-and-off cycles of UV illumination **(Figure S4e)**, with high photoresponsivity indicated by shorter rise (τ_rise_ = 0.35 s) and decay times (τ_decay_ = 0.74 s) compared to PduA **(Figure S4fi-ii)**. These experiments suggest that PCET is the primary mechanism underlying photocurrent generation in shell proteins. Overall, the enhanced electron conduction and photocurrent generation observed in the PduBB’ is driven by its higher number of tyrosine residues. We examined the energy level difference in the shell proteins to gain further insights into electron conduction.

To explore the energy levels in the shell proteins, we prepared an energy band diagram by analyzing the Ultraviolet Photoelectron Spectroscopy (UPS) data, complemented by the Tauc plot derived from the absorbance spectra of the proteins **(Figure 4, Figure S3)**.

**Figure 4:**
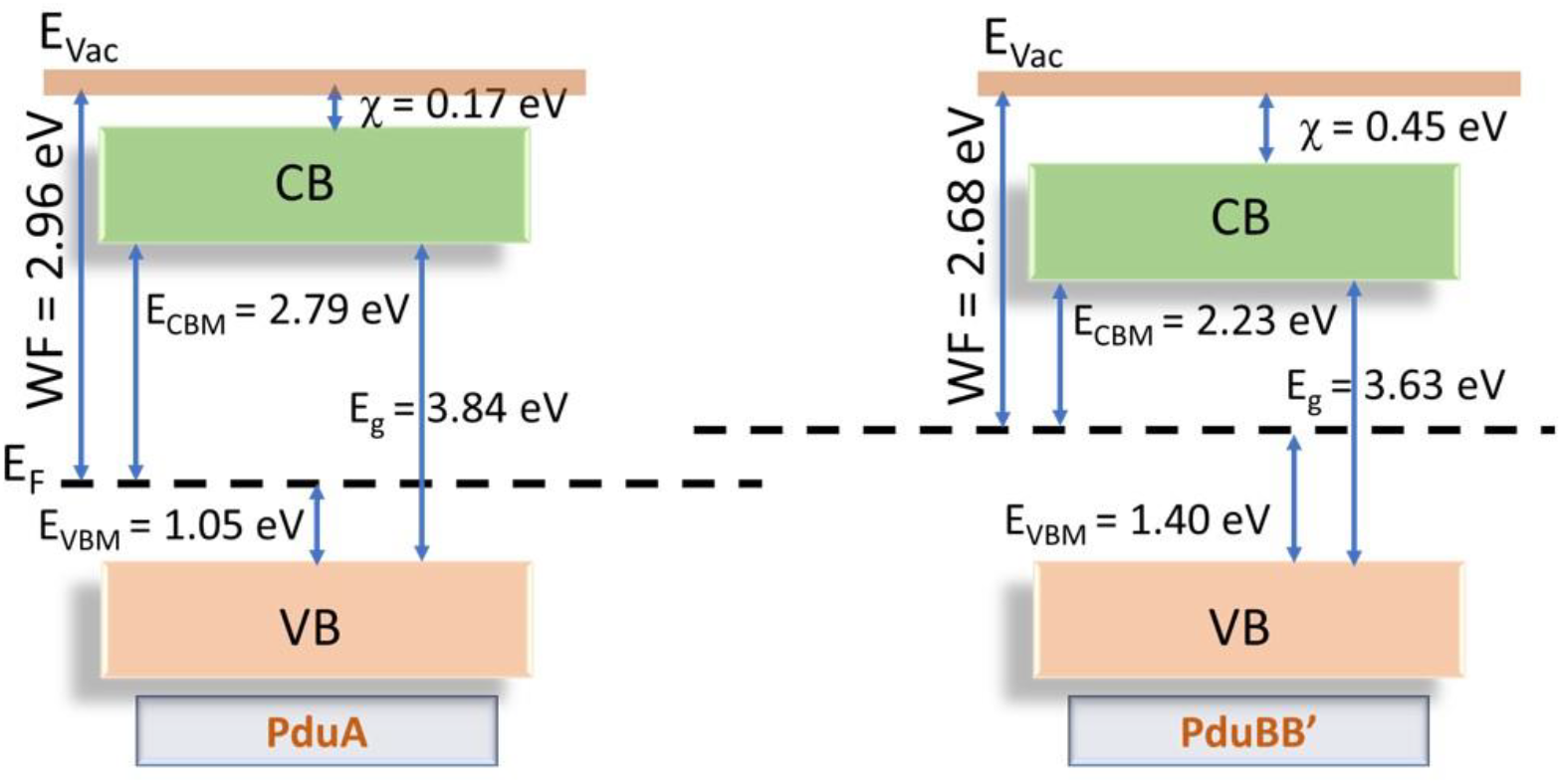
Energy band diagram of shell protein PduA and PduBB’: A plausible energy band diagram prepared using data of work function calculated from UPS data complemented with Tauc plot derived from the absorbance spectrum of shell proteins.

The work functions determined from UPS were 2.96 eV for PduA and 2.63 eV for PduBB’. Additionally, we analyzed the valence band maximum (VBM) positions, extrapolating them from the UPS spectra. The estimated VBM positions by (regarding Binding Energy) are 1.05 and 1.40 eV for PduA and PduBB’, respectively. When combined with the respective band gaps of 3.84 eV (PduA) and 3.63 eV (PduBB’), we observed a significant shift in the Fermi level toward the conduction band minimum (CBM), particularly for PduBB’.

The VBM positions further indicate a substantial number of unoccupied states in the conduction band, facilitating electron mobility and higher conductivity, particularly for PduBB’. The deeper VBM position of PduBB’ relative to its Fermi level indicate a large energy gap between the filled valence band and the empty conduction band. This gap minimizes electron trapping in the valence band, allowing them to efficiently transition into the conduction band, contributing to increased electron flow and enhanced current conduction. This mechanism could explain the superior photocurrent and corresponding performance metrics observed in the self-assembled PduBB’ compared to PduA.

The band gaps of the shell proteins PduA (3.84 eV) and PduBB’ (3.63 eV) position them within the range of well-known wide band gap semiconductors, such as gallium nitride (GaN, ∼3.4 eV) and silicon carbide (SiC, ∼3.2 eV). These materials are widely used in high-power electronics, UV photodetectors, and high-temperature applications due to their ability to handle high voltages and perform reliably under extreme conditions. However, they come with limitations of high fabrication costs, the need for complex epitaxial growth techniques, and integration challenges with flexible or biocompatible materials^37,38^. In this regard, the shell proteins offer several advantages that address these limitations. Their natural propensity to self-assemble into nanostructures like nanosheets and nanotubes provides a cost-effective, bottom-up fabrication method, eliminating the need for expensive fabrication processes. Furthermore, as biological materials, these proteins are inherently flexible and biocompatible, making them ideal for soft electronics, wearable devices, or bio-hybrid systems. Their electronic properties can be tailored through natural or engineered modifications, offering enhanced versatility for developing customizable semiconducting materials. We further evaluate the performance of shell proteins as photodetectors, examining their response characteristics.

The semiconductive and photoresponsive properties of the shell proteins make them ideal candidates for photodetector applications. A crucial factor in assessing a photodetector is its sensitivity towards measured the photoresponse of both PduA and PduBB’ shell proteins under varying UV light power (0.11 mW, 0.15 mW, 0.25 mW, and 0.31 mW), as shown in **(Figures 5a and 5b)**. We observed a corresponding increase in photocurrent generation with increasing UV light power.

**Figure 5:**
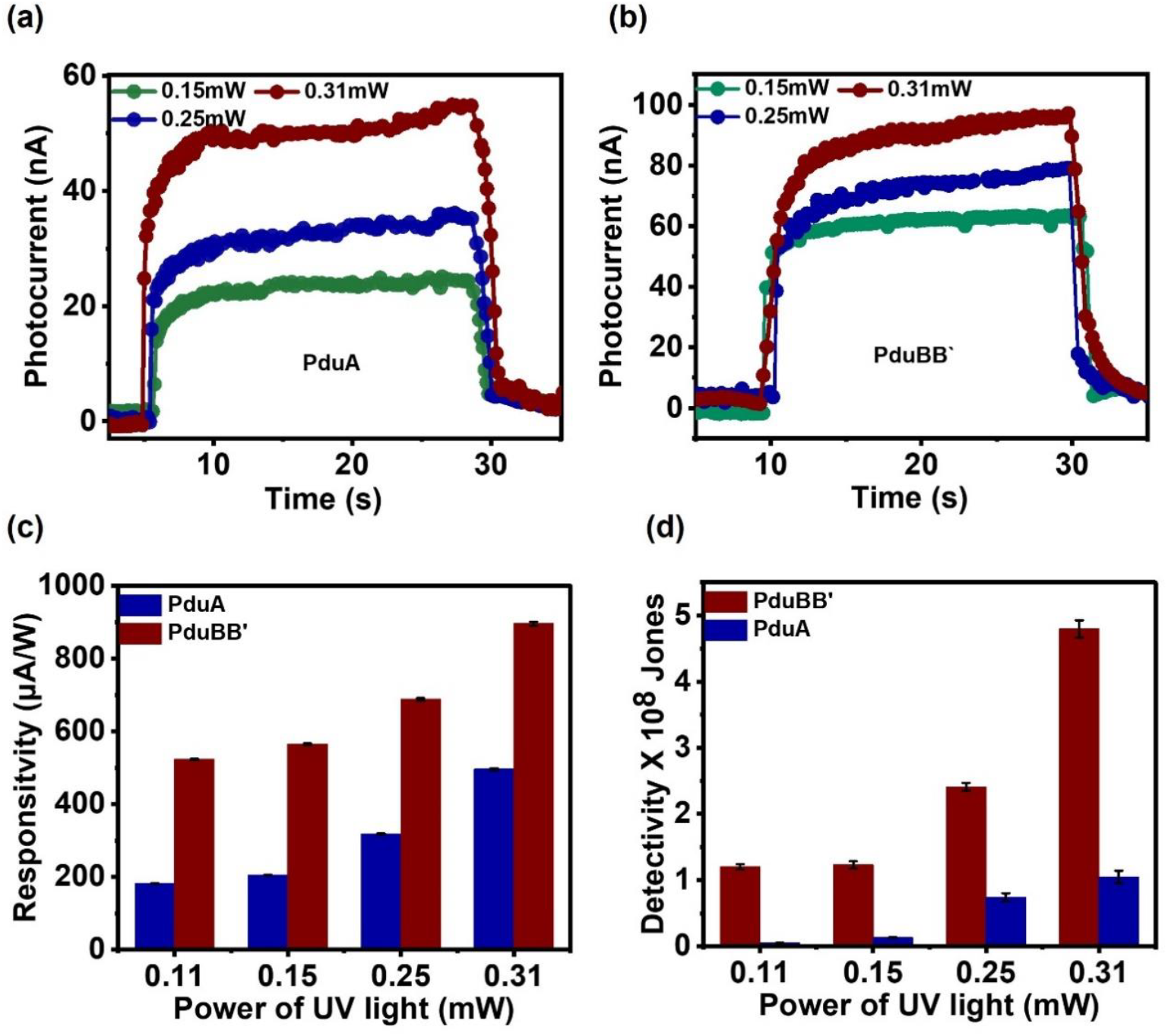
Photodetector performance metrics of PduA and PduBB’ shell proteins: (a, b) Power-dependent photoresponse of PduA and PduBB’, illustrating increased photocurrent generation with rising UV light power, highlighting the sensitivity of both proteins to UV illumination; (c) Responsivity of PduA and PduBB’ to incident UV light, demonstrating rapid current generation upon illumination, indicating their potential for fast photoresponse applications; (d) Detection limits of PduA and PduBB’ under varying UV light power (0.11 mW, 0.15 mW, 0.25 mW, and 0.31 mW), as depicted in Figures 5a and 5b, showcasing the capability of these shell proteins to detect low-power light and function efficiently across a range of illumination conditions.

The PduA shell protein exhibited an increase of photocurrent from 20 nA to 50 nA, while PduBB’ showed a surge from 50 nA to 90 nA. This highlights the significant potential of shell proteins to be integrated into electronic devices, enabling the development of efficient bio-photodetectors.

To further evaluate the photodetector performance of the shell proteins, we estimated key figures of merit, including responsivity (R), detectivity (D*), and external quantum efficiency (EQE)^39^ as shown in **(Figures 5c, 5d, and Table 1)**, using equations 1-3 described in the materials and methods section.

**Table 1:** The calculated enhanced quantum efficiency for PduA, PduBB’ at varying UV light power from 0.11-0.31mW, indicating high efficiency of PduBB’ to convert incident photons to electrons.

| UV light Power | PduA (EQE %) | PduBB' (EQE %) |
| --- | --- | --- |
| 0.11 mW | 0.1% | 0.25% |
| 0.15 mW | 0.11% | 0.28% |
| 0.25 mW | 0.15% | 0.35% |
| 0.31 mW | 0.24% | 0.43% |

The PduBB’ shell protein demonstrated higher values across all figures of merit compared to PduA, Figure 5c, 5d, and Table 1, indicating higher sensitivity and superior photodetector performance. This enhanced performance is likely attributed to the spatial arrangement and increase in tyrosine residues in PduBB’. These parameters provide critical insights into the photoresponse behavior and sensitivity of the shell proteins within our microcompartment system.

In summary, the shell proteins, particularly PduBB’, exhibit exceptional photodetector performance with high responsivity, detectivity, and external quantum efficiency. Their superior sensitivity and efficiency compared to existing protein-based photodetectors underscore their potential for advanced optoelectronic applications.

In this study, we investigated the potential of shell proteins, specifically PduA and PduBB’, as advanced photodetectors for UV light detection. The strategic arrangement of tyrosine residues in the shell proteins results in photocurrent generation and overall sensitivity, underscoring the pivotal role of tyrosine-induced proton-coupled electron transfer (PCET) in efficient UV light detection. Our results highlighted the exceptional semiconducting and photoresponsive properties of shell proteins, with PduBB’ demonstrating superior performance over PduA.

Shell proteins exhibit high responsivity, detectivity, and external quantum efficiency, making them promising candidates for advanced bio-photodetectors. PduBB’ demonstrates a threefold increase in photocurrent compared to PduA and maintains high photoresponsivity across extended UV illumination cycles. These performance advantages are attributed to the unique spatial arrangement of tyrosine residues in PduBB’, which significantly boosts the photodetector’s capabilities.

Compared to existing protein-based photodetectors and UV-enhanced systems, our shell proteins offer superior performance metrics, including notably higher EQE (**Table 2**). Unlike previously reported protein functionalized systems, our bacterial microcompartment shell proteins leverage their inherent distinct structural and electronic properties for photoresponsive applications. These unique characteristics position shell proteins as a promising choice for next-generation photodetectors, surpassing the performance of traditional protein-based systems.

**Table 2:** compares the external quantum efficiency (EQE) of our PduBB’ shell protein to that of a photosynthetic protein coupled with UV enhancer molecules, revealing that PduBB’ achieves a remarkable EQE of 0.43%, substantially higher than the 0.048% EQE of the reported protein system^40^.

| Protein System | EQE % |
| --- | --- |
| Photosynthetic protein coupled with UV enhancer molecules | 0.048% |
| Shell protein PduBB' | 0.43% |

Looking forward, we plan to explore the integration of these proteins with functional moieties emitting in the visible range of light to extend their detection range from UV to visible light. Future research may focus on optimizing shell proteins in functional devices, evaluating their photodetection performance under various environmental conditions, and scalability for broader applications. The versatility and enhanced sensitivity of shell proteins offer exciting opportunities for developing advanced photodetectors and their integration into flexible and wearable electronics, paving the way for innovative technologies.

## Author Contributions

SB: conceptualization, experimental design, execution, methodology, data analysis, investigation, manuscript drafting, and editing. SMR: experimental execution, data analysis, manuscript writing, and editing. SS: Supervision, project investigation, conceptualization, design, analysis, writing, reviewing, and editing.

## Acknowledgments

SB acknowledges DST-INSPIRE for fellowship and INST for research facilities. SMR acknowledges INST for fellowship and facilities. The authors also acknowledge all the SS Lab members for insightful discussions.

