## supplementary files for "Illuminating the Potential: Light Harvesting and Semiconducting Properties of Bacterial Microcompartment Shell Proteins"

**Experimental Section**

Unless otherwise mentioned, all the chemicals were procured from Sigma-Aldrich (India). Ultrapure water was used throughout the experiments. pET41a vector is a kind gift from Prof. T.A. Bobik, Iowa State University, U.S.A.

**Cloning, expression, and purification of PduBMC shell proteins**

To mutate tyrosine to alanine (PduA Y35A) and tryptophan (PduA Y35W) of PduA shell protein, we used overlap extension PCR using primers mentioned in Tables S1 and S2. The point mutation was confirmed by Sanger sequencing. The mutant PCR product with BglII and HindIII restriction sites was cloned into a pET41a vector having a kanamycin-selectable marker and transformed into chemically competent *E. coli* DH5α cells. The positive colonies were selected on kanamycin plates. The recombinant plasmids were purified and transformed into BL21(DE3) cells for protein expression.

To purify the shell proteins, 1% of overnight grown culture was inoculated in bulk secondary media and incubated for 1.5 to 2 hours until it reached an $OD_{600}$ of 0.5. The expression was induced by adding 0.5 mM IPTG to the secondary culture and incubated at 28°C for 12 hours. The cells were harvested by centrifugation and lysed in lysis buffer (50 mM Tris-base pH 7.5, 200 mM NaCl, and 5 mM imidazole) along with lysozyme (2.3 mg/g of cell pellet) and PMSF (0.5 mM). Further, cells were sonicated and centrifuged at 11000 rpm for 30 minutes. The supernatant was loaded to the Ni-NTA affinity column, pre-equilibrated with column buffer, and washed with wash buffer (50 mM Tris Base, 200 mM NaCl, and 50 mM imidazole; pH 7.5). The bound protein was eluted using elution buffer (50 mM Tris Base, 200 mM NaCl, and 200 mM imidazole; pH 7.5). The eluted fractions were checked for purity on SDS-PAGE. The concentration of the protein was estimated by Bradford assay.

### TEM imaging

The individual protein samples of the shell protein PduA and PduBB` were prepared by drop casting 10 µl (5 µM) of the protein on the formvar-coated 300-carbon mesh grid followed by adding 10 µl of 1% uranyl acetate for negative staining. Samples were incubated for 30 s in the dark, followed by washing with water and air drying. TEM imaging of the protein samples was done by using JEM 2100 TEM (JEOL, USA) operated at 120kV.

### Field-emission scanning electron microscopy (FESEM) Imaging:

The morphology of PduA and PduBB` is further confirmed by performing FESEM imaging. The silicon wafers were cleaned with distilled water followed by isopropyl alcohol. The 10µl of 5 µM protein concentration of both the shell proteins is dropcasted on the cleaned and dried Silicon wafer. The sample was allowed to incubate on the wafer for 3 minutes followed by wicking off with the Whatman Filter paper. The sample was allowed to dry in air followed by vacuuming in a desiccator. The imaging of the sample has been done using the FESEM instrument, Model Number: JEOL JSMIT 300.

### Ultraviolet photoelectron spectroscopy

Shell proteins were dropcasted on the cleaned ITO substrate and allow for air-drying overnight. The ITO substrate was thoroughly cleaned following established protocols before drop-casting the protein solution onto it, ensuring the formation of a uniform film. The UPS spectra were recorded using a pass energy of 2 eV and a step size of 0.05 eV, with a He I source (hν = 21.22 eV). The analyser lens axis was set at 90˚ relative to the sample surface, and the take-off angle was fixed at approximately 30˚ to focus on a smaller solid angle. The instrument was calibrated using a gold target sample in an ultrahigh vacuum chamber, with a spectrometer resolution of 0.12 eV, to ensure accuracy. Data analysis focuses on determining the work function of proteins.

### I-V characterization

We used an ITO-coated glass substrate ($1 \times 1$ $cm^2$) for I-V characterization. First, we cleaned the substrate with IPA and acetone solution followed by bath sonication for 5 minutes. The substrate was dried and a channel of a few microns wider was created over the ITO substrate using a diamond cutter. Two external copper wires were attached on both sides of the electrode for electrical measurement. Subsequently, the sample solution was cast upon the channel to make a proper interface between two sides of the ITO electrodes. Further, voltage bias has been applied to the sample to measure the change in the current using a source meter.

**Microscopic imaging of ITO substrate**

The channel cast on the ITO substrate is imaged under a 10X objective lens using a microscop Olympus lX73. Images were further analyzed using ImageJ software to calculate the width and length of the casted channel.

**Photocurrent experiment**

To check the photo response of the material, we irradiate the sample using short UV light of 254 nm periodically (on and off) and check the change in the current concerning the light under non-biased conditions.

We calculated the $\tau_{rise}$ when the light source is turned on and it reaches the maximum photocurrent. Similarly, for $\tau_{decay,}$ the difference between the maximum current and the lower optimum photocurrent is taken on switching off the light. The area used to calculate both times is represented in from the zoomed area shown in Figures 2f and 4f. Further, to interpret the photodetector's performance, we have estimated various figures of merit such as responsivity (R), detectivity (D*), and external quantum efficiency (EQE), summarized in Table 1, using the following standard equations[39]

$$R = \frac{I_P}{P^* \times A} \quad \text{.........................} (1)$$

$$D^* = \frac{R\sqrt{A}}{\sqrt{2qI_d}} \quad \text{.........................} (2)$$

$$EQE = \frac{h.c.R}{q\lambda} \quad \text{.....................} (3)$$

where, "Ip" refers to the **photocurrent**, which was the current generated in the presence of UV light illumination, while "Id" denotes the **dark current**, which is the baseline current measured in the absence of light. $P^*, q$, $A$, and $\lambda$ are the incident light illumination power density, elementary charge, effective area of PD, and incident wavelength, respectively. Responsivity quantifies how efficiently the photodetector converts incoming light into an electrical signal, measured as the ratio of photocurrent (I) to the incident optical power (P) on the active area.

Detectivity represents the sensitivity of the photodetector to weak light signals, while EQE reflects the efficiency of converting incident photons into electrons, defined as the number of charge carriers generated per photon[39].

**Tabel S1:**

**Protein Sequence of PduA:**

MQQEALGMVETKGLTAAIEAADAMVKSANVMLVGYEKIGSGLVTVIVRGDVGAVK
AATDAGAAAARNVGEVKAVHVIPRPHTDVEKILPKGISQ

**Protein Sequence of PduBB`:**

MSSNELVEQIMAQVIARVATPEQQAIPGQPQPIRETAMAEKSCSLTEFVGTAIGDTLGL
VIANVDTALLDAMKLEKRYRSIGILGARTGAGPHIMAADEAVKATNTEVVSIELPRDT
KGGAGHGSLIILGGNDVSDVKRGIEVALKELDRTFGDVYGNEAGHIELQYTARASYA
LEKAFGAPIGRACGIIVGAPASVGVLMADTALKSANVEVVAYSSPAHGTSFSNEAILVI
SGDSGAVRQAVTSAREIGKTVLATLGSEPKNDRPSYI

| Protein | No. of Tyrosine |
|---|---|
| PduA | 1 |
| PduBB’ | 6 |

**Table S1:** Protein sequences of PduA and PduBB` representing the number of tyrosine residues highlighted in red. PduA has one tyrosine residue instead PduBB` is having 6 residues per monomeric unit.

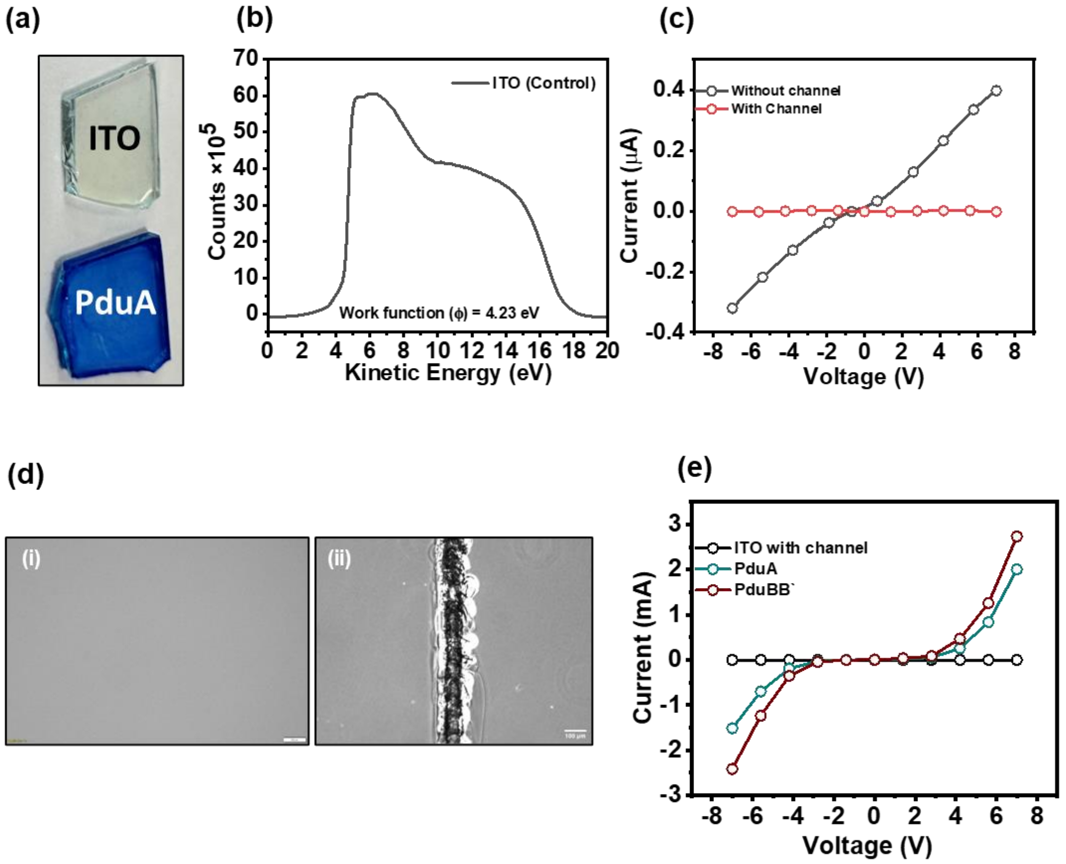


**Figure S1: Electrical conductivity of control ITO substrate:** (a) represents the bare ITO substrate and protein thick layer cast over the ITO substrate, stained with Coomassie Brilliant Blue R-250 dye to ensure protein coverage over the substrate; (b)Measured work function value of control ITO substrate; (c) I-V response of bare ITO without channel showing a linear increase in current on applying bias voltage corresponding to its conducting behavior and disruption in conductivity on casting channel on ITO substrate; (d) optical microscopic imaging of bare ITO substrate and casted channel over the ITO substrate surface, Scale bar- 100µM; (e) I-V response of ITO substrate with bare channel showing no electron conduction and in presence of protein sample that bridges the connection between two ends of ITO substrate and show flow of current in a non-linear manner.

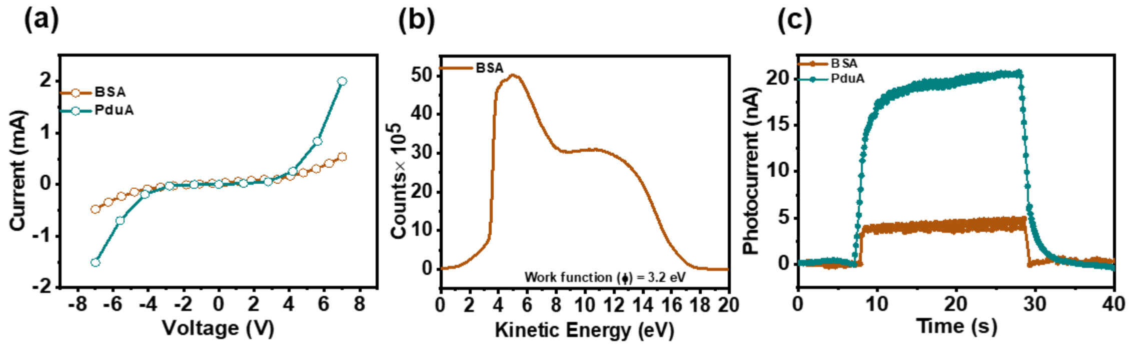


**Figure S2: Electrical and photoresponsive properties of BSA:** (a) I-V curves for globular protein BSA and shell protein PduAWT, demonstrating higher current flow for PduAWT compared to BSA; (b) Calculated work function (ϕ) values for BSA and shell proteins, with BSA showing a higher work function relative to shell proteins; (c) Photocurrent responses of BSA and PduAWT under UV light (254 nm) illumination, indicating that shell proteins exhibit significantly higher photo-responsivity than BSA.

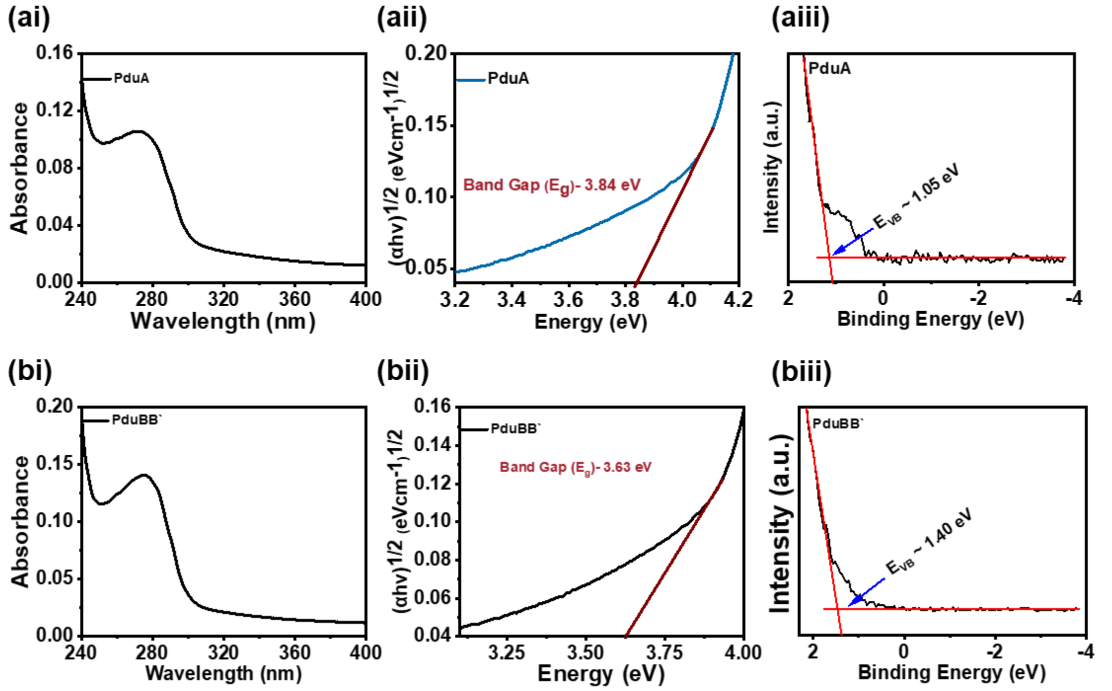


**Figure S3: Band gap calculation for shell protein sheets:** (ai) Absorbance spectrum of PduA (aii) Tauc plot derived from the absorbance spectrum to calculate the band gap for PduA shell protein; (aiii) Valence band calculation by extrapolating the UPS spectrum of PduA protein; (b i-iii) Absorbance spectrum, Tauc plot and extrapolated UPS spectrum for PduBB` shell protein.

| | Primers used to generate mutant PduAY35A |
|---|---|
| **Forward Primer (BglII)** | Gtgatgttagtgggcgccgaaaagattggc |
| **Reverse Primer (HindIII)** | Gccaatcttttcggcgcccactaacatcac |
| **Confirmed Sequence** | **atg**caacaagaagcactaggaatggtagaaaccaaaggcttaaccgcagccatagaggccgctgatgcaatggttaagtcagccaat[illegible]<br>ggcgccgaaaagattggctccgggctggtaaccgtcatcgtgcgcggcgatgttggcgcggtcaaagcggccaccgatgcaggtgcc[illegible]<br>gcaacgtgggtgaagtgaaagccgtacacgtcatcccacgccctcacaccgatgtagaaaaaatcttaccgaagggaattagc[illegible] |

**Table S2:** Forward and reverse primes create a point mutation in PduAWT to mutate tyrosine present at 35th position to alanine, generating a PduAY35A mutant. Point mutation is confirmed by doing the Sanger sequencing of the cloned plasmid having the gene of interest.

| | Primers used to generate mutant PduAY35W |
|---|---|
| **Forward Primer (BglII)** | 5'-gcagcaagatctatgcatcaccatcatcaccaccaacaagaagcacta-3' |
| **Reverse Primer (HindIII)** | 5'-gcctgcaagctttcattggctaattcc-3' |
| **Confirmed Sequence** | **atg**caacaagaagcactaggaatggtagaaaccaaaggcttaaccgcagccatagaggccgctgatgcaatggttaagtcagccaatgt<br>gatgttagtgggcccagaaaagattggctccgggctggtaaccgtcatcgtgcgcggcgatgttggcgcggtcaaagcggccaccgatgca<br>ggtgccgcagccgcacgcaacgtgggtgaagtgaaagccgtacacgtcatcccacgccctcacaccgatgtagaaaaaatcttaccgaa<br>gggaattagcca**atga** |

**Table S3:** Forward and reverse primes used to create point mutation in PduAWT to mutate tyrosine present at 35$^{th}$ position to try, generating PduAY35W mutant. Point mutation is confirmed by doing the Sanger sequencing of the cloned plasmid having the gene of interest.

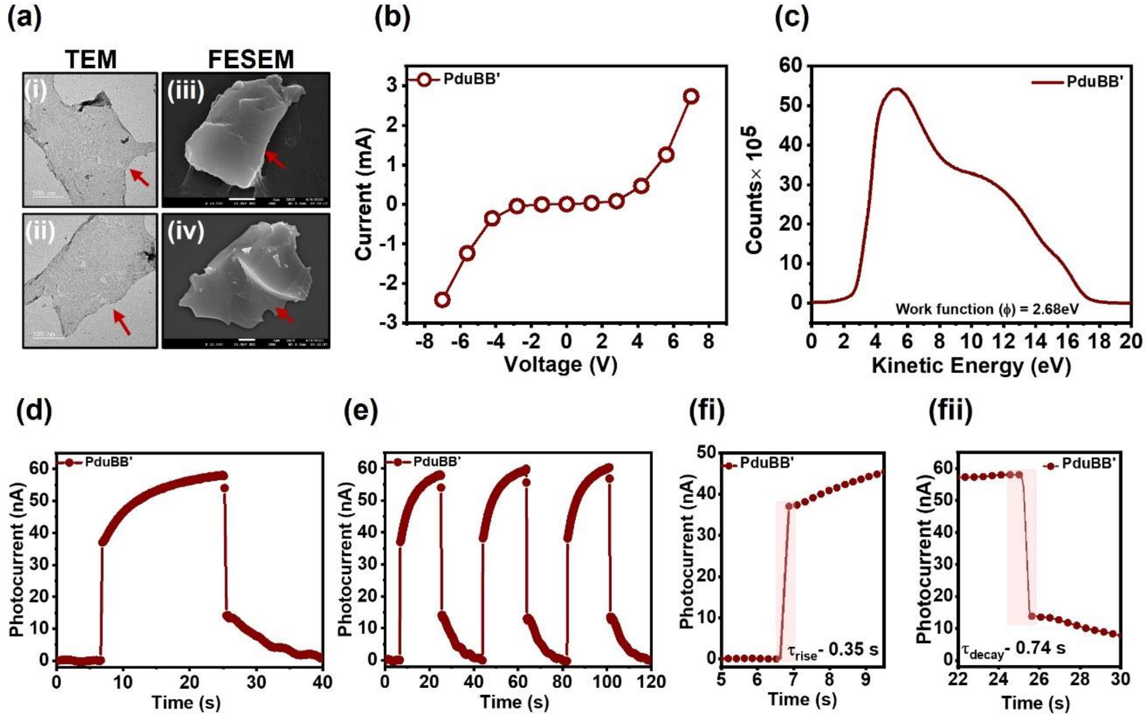


**Figure S4: Electron Conductivity and Photocurrent Generation of PduBB` Shell Protein:** (a)TEM and FESEM imaging of PduBB` showing sheet forming tendency of the shell protein; (b) I-V characteristics of PduBB`, showing a higher current flow compared to PduA, suggesting enhanced electron conductivity in the PduBB` shell protein; (c) Photocurrent generation observed in PduBB` under UV light illumination (254 nm), demonstrating its capacity for light-driven electron transport; (d) on-off cycles of photocurrent response under alternating dark and UV light conditions, showing stable and reversible photocurrent generation; (e) $\tau_{rise}$ and $\tau_{decay}$ time profiles for the PduBB` photoresponse, indicating high light sensitivity and fast response times, suggesting PduBB` as a promising candidate for light-harvesting application
